# Nanoparticle-enabled plasma proteomics of a mouse atherosclerosis model

**DOI:** 10.1101/2025.08.01.667173

**Authors:** Constance Delwarde, Joan T. Matamalas, Sarvesh Chelvanambi, Taku Kasai, Gabriel Shlayen, Diego V. Santinelli-Pestana, Maedeh Zamani, Elena Aikawa, Masanori Aikawa, Sasha A. Singh

## Abstract

**Background:** Dyslipidemia, marked by elevated LDL-cholesterol (LDL-C), is a major risk factor for coronary heart disease. Mouse experimental models, such as *Ldlr^-/-^* mice that develop atherosclerosis and metabolic disorders when fed a high-fat diet (HFD), are indispensable for studying disease mechanisms and identifying potential biomarkers.

**Objectives:** We aimed to profile the plasma proteins in experimental murine atherosclerosis with the primary goal of detecting low-abundant proteins.

**Methods:** *Ldlr^-/-^* mice were fed a chow diet or HFD for 3 (n = 27 per group) or 6 months (n = 12 per group). Plasma samples were processed using nanoparticle technology (Proteograph^®^ XT Assay; Seer, Inc), and peptides were analyzed using the Orbitrap Astral (Thermo Fisher Scientific) in data-independent acquisition mode. For tissue proteomics, ten aortas were pooled for n = 4 pools per time point and diet, and n = 6 livers per time point and diet. Peptides were analyzed on the Orbitrap Exploris 480 in data-dependent mode. Proteomes were queried against the Tabula Muris mouse single-cell, STRING, and Gene Ontology databases, and queried against a genome-wide association list of 419 risk loci for coronary artery disease.

**Results:** We sequenced 5,080 plasma proteins, surpassing previous reports by 10-fold. The prototypical apolipoproteins and complement factors were the most abundant proteins, whereas proteins associated with cytokine/chemokine signaling represented the previously uncharted mouse plasma proteome. We divided the proteome into quartiles (Q1-Q4) to monitor sweeping changes over time. Proteins with a sustained enrichment in HFD (n = 705) were indicative of liver cell subtypes (Tabula Muris). Whereas proteins that moved up from the lower quartiles - Q2 (n = 228), Q3 (115) and Q4 (63) – associated with leukocyte, fibroblasts, and endothelial cell markers, indicating that signatures of inflammation and endothelial activation increase with disease progression. Notably, 86 and 146 proteins were increased at 3 and 6 months, including MMP-12 and COL6A3. Classical apolipoproteins exhibited heterogeneous responses - SAA3 and APOC2 increased, while APOA1, APOE, and LCAT decreased with high-fat feeding, suggesting impaired high-density lipoprotein (HDL) functionality. Proteins shared between plasma and aorta were enriched for extracellular matrix components, while those overlapping with liver reflected metabolic processes. Finally, 120 coronary artery disease (CAD)-associated proteins from human GWAS were detected in *Ldlr^-/-^* plasma, of which 4, including lipoprotein lipase, exhibited an increase in abundance with HFD.

**Conclusions:** Nanoparticle-dependent proteome enrichment coupled to mass spectrometry may allow to identify novel plasma biomarkers in *Ldlr^-/-^* mice and facilitate monitoring of candidate proteins associated with human disease mechanisms in preclinical interventional studies, thereby opening new avenues for understanding disease pathology and uncovering novel molecular contributors.

## INTRODUCTION

Atherosclerosis remains one of the leading causes of morbidity and mortality worldwide, contributing significantly to the global burden of cardiovascular disease.^1^ The pathogenesis of atherosclerotic vascular disease is complex and multifactorial, involving lipid metabolism disturbances and chronic inflammation.

Dyslipidemia promotes atherosclerosis primarily by elevated plasma levels of low-density lipoprotein-cholesterol (LDL-C).^2–6^ Atherosclerosis is a chronic disease whose progression cannot be easily studied in humans; therefore, animal models, particularly mouse models, are essential for research.^7, 8^ The two most common mouse models for *in vivo* atherosclerosis studies rely on genetic deletion of either *Apoe* or *Lldr.*^9–11^ The smaller high-density lipoproteins (HDL) usually predominate the plasma of wild-type mice, but in *Apoe^-/-^* and *Ldlr^-/-^* mice, plasma lipoprotein species shift towards the APOB100/48 particles, VLDL, IDL, and LDL. *Apoe^-/-^* mice accumulate mostly VLDL and IDL, whereas *Ldlr^-/-^* mice accumulate VLDL and LDL in a high-cholesterol, high-fat diet.^9–13^ LDL deposition into the subendothelial space is one of the earliest events in atherosclerosis, which precedes immune responses, such as macrophage accumulation in humans.^14^ LDL’s retention within the intima is facilitated by binding to extracellular matrix components such as proteoglycans and collagen.^14, 15^

To identify candidate disease drivers of atherosclerosis in the *Ldlr^-/-^*model, molecular profiling workflows such as proteomics have examined solid organs including the aorta and liver.^16–19^ Such studies aim to characterize disease progression driven by local and remote (organ-to-organ crosstalk) processes.^16^ Although circulating proteins are expected to reflect key disease mechanisms and serve as potential biomarkers, the murine plasma proteome has remained an underexplored resource in cardiovascular disease research. Plasma/serum is dominated by approximately 300 proteins, including albumin, apolipoproteins, immunoglobulins, fibrinogens, and complement factors, that mask the presence of potential novel disease-informative biomarkers that span at least a 10^10^-fold dynamic range.^20, 21^ This masking phenomenon has been particularly challenging for mass spectrometry-dependent profiling. However, multiple gains have been made on the human plasma front, primarily to leverage the common practice of acquiring body fluids to monitor biomarkers with prognostic or diagnostic value. These and other clinical applications have sustained interest in innovative deep plasma proteome profiling.^21^ Affinity binding-dependent workflows (using aptamers or antibodies) emerged over a decade ago. They rely on an amplicon or fluorophore readout for protein detection.^22^ These approaches provide panels surpassing 4,000 target human proteins, but limitations of these technologies exist, including the lack of non-human panels, which limits their applicability in animal studies.^22^

Recent advances in nanoparticle-dependent protein corona-forming workflows have made mass spectrometry plasma proteomics increasingly feasible.^23^ The engineered surfaces of these nanoparticles bind proteins in two phases - first non-specifically by high-abundant proteins, followed by high-affinity lower-abundant proteins that displace the former at the surfaces of the nanoparticles; a processes first described by Vroman and colleagues.^24^ The nanoparticle enrichment method therefore binds proteins based on chemistry rather than abundance,^25^ thereby enriching lower abundant proteins and compressing the dynamic range of protein signal for the mass spectrometer. Moreover, mass spectrometers have become increasingly faster, achieving more than 5,000 proteins within a 30-minute runtime, which is markedly quicker than conventional times of 60 minutes or more to yield comparable results. These platforms provide researchers with the means to implement mass spectrometry-based workflows to profile hundreds of preclinical or clinical plasma samples in a matter of days.^26^ We therefore took advantage of a nanoparticle-enrichment strategy previously applied to human plasma,^27, 28^ to profile the plasma proteome of *Ldlr^-/-^* mice fed a HFD or chow diet for 3 or 6 months, timepoints during which plaques form and become progressively diseased.^13, 18, 29, 30^

## METHODS

### Animal experiments

All animal procedures were approved and performed in compliance with Beth Israel Deaconess Medical Center’s Institutional Animal Care and Use Committee (IACUC Protocol # 003-2020). Eight-week-old male *Ldlr^-/-^*mice were acquired from the Jackson Laboratory (RRID:IMSR JAX:002207). Upon arrival, the mice immediately began either a chow/control diet (LabDiet, catalog #5008i) or HFDs (Clinton/Cybulsky High Fat Rodent Diet (1.25% Chol), gamma-Irradiation; Research Diets, Inc., catalog #D12108Ci) for 3 (n = 27) or 6 (n = 12) months.

### Plasma sample preparation with the Proteograph^®^ XT Assay protocol

Mice were euthanized, using the CO^2^ chamber. Then, blood was collected from the left ventricle.^31^ Plasma samples were processed with Proteograph^®^ Product Suite using the Proteograph XT Assay,^32^ using 75 µl of each plasma sample that was mixed with 165 µl of Reconstitution Buffer A (S55R3090) to yield a final volume of 240 µl and were then transferred to Seer Sample Tubes for automated processing with the Proteograph XT Assay kit (S55R1100). Plasma samples were aliquoted into two wells of a Sample Prep Plate, and proteins were quantitatively captured in protein coronas associated with nanoparticle suspensions A and B. Proteins were subsequently denatured, reduced, alkylated and subjected to proteolytic digestion (trypsin and Lys-C). Peptides were purified and yields were determined (Thermo Fisher Scientific, catalog #23290). Peptides were dried down overnight with a vacuum concentrator and reconstituted with a reconstitution buffer to a concentration of 50 ng/µl. All plasma samples in the study were processed across two Proteograph^®^ XT Assay plates.

### Plasma sample preparation with the direct digestion method

For direct digestion of 5 µl diluted neat plasma samples, proteins were denatured, reduced, alkylated, and subjected to proteolytic digestion (Trypsin and Lys-C) for 3 hours at 37°C. Peptides were purified by solid phase extraction, and yields were determined (Thermo Fisher Scientific catalog #23290).

### Data-independent acquisition (DIA) mass spectrometry

For data-independent acquisition (DIA), 8 µl of reconstituted peptide mixture from each nanoparticle preparation was analyzed, resulting in a constant 400 ng load on column for nanoparticle A and nanoparticle B samples. Each sample was analyzed with a Vanquish NEO nanoLC system coupled with an Orbitrap ^TM^ Astral ^TM^ (Thermo Fisher Scientific) mass spectrometer using a trap-and-elute configuration. First, the peptides were loaded onto an Acclaim^TM^ PepMap^TM^ 100 C18 (0.3 mm ID x 5 mm) trap column and then separated on a Gen1 50 cm µPAC^TM^ analytical column (Thermo Fisher Scientific) at a flow rate of 1 µl/min using a gradient of 4 – 35% mobile phase B (acetonitrile/0.1% formic acid) mixed into mobile phase A (water/0.1% formic acid) over 21 minutes, resulting in a 26 minute run time. The mass spectrometer was operated in DIA mode. MS1 scans were performed in the Orbitrap detector at 240,000 resolution, every 0.6 seconds with a 5 ms ion injection time or 500% automatic gain control (AGC; 500,000 ion) target. For DIA experiment, two hundred fixed DIA window covering a 380 Th – 980 Th mass range were collected at the Astral detector per cycle with 3 Th precursor isolation windows, 25% normalized collision energy, and 5 ms ion injection times with a 500% (50,000 ions) active gain control maximum. MS2 scans were collected from m/z 150-2000. Window placement optimization was turned on. A source voltage of 1500 V and an ion transfer tube temperature of 300°C were used for all experiments.

### MS/MS spectral annotation for the plasma proteomes

DIA data was processed using Proteograph^®^ Analysis Suite (PAS). The spectra were processed using the DIA-NN search engine^33^ (version 1.8.1) in library-free mode searching MS/MS spectra against an *in silico* generated spectral library based on the Uniprot mouse database (UP000000589_10090), excluding fragment sequences, resulting in 46,734 mouse entries and an additional 91 contaminant entries,. Library-free search parameters include trypsin protease, 1 missed cleavage, N-terminal methionine excision, fixed modification of cysteine carbamidomethylation, peptide length of 7-30 amino acids, precursor range of m/z 300-1800, and fragment ion range of m/z 200-1800. MS1 and MS2 mass accuracy was set to 3 and 8 ppm, respectively. Precursor and Protein Group false-discovery rate (FDR) thresholds were set at 1%. Quantification was performed on summed abundances of all unique peptides considering only precursors passing the FDR thresholds. PAS summarizes all nanoparticle values for a single protein into a single quantitative value. Specifically, a single protein may have been measured two times, once for each nanoparticle. To derive the single measurement value, PAS uses a maximum representation approach, whereby the single quantification value for a particular peptide or protein group represents the quantitation value of the nanoparticle most frequently measured across all samples.^34^

### Liver and aorta homogenization

The protocol was based on our previously published methods.^35^ Before the standard homogenization step, each frozen aorta was minced. Each tissue was transferred to a Precellys 2 ml tube from the Soft Tissue Homogenizing Ceramic Beads Kit (Bertin Technologies, catalog #CK14) for liver, and from the Hard Tissue Homogenizing Ceramic Beads kit (Bertin Technologies, t catalog #CK28) for aorta. Then, the same protocol was applied to the aorta and liver frozen samples. A modified RIPA buffer [0.5 ml, 50 mM Tris-HCl pH 7.4 (Boston BioProducts, catalog #BM-327), 0.4 M NaCl (Sigma-Aldrich, catalog #S9888), 1.0 mM EDTA (Boston Bio Products, catalog t#BM-150), 1.0% nonidet P-40 (Sigma-Aldrich, catalog #74385), 0.1% sodium deoxycholate (Sigma-Aldrich, catalog #D6750), 40 µM PJ34, 1.0 µM ADP-HPD, protease inhibitor cocktail (Sigma-Aldrich, catalog #P8340), phosphatase inhibitor (Sigma-Aldrich, catalog #4906845001)] was added to tubes for protein analysis. Tissues were homogenized in a Precellys 24 Tissue Homogenizer (Bertin Technologies, catalog #P000669-PR240-A) using three 10-second cycles at 5,000 rpm that were then cooled on ice for 15 minutes. Tissue debris was then removed by centrifugation (3,000 rpm, 4 °C, 5 min).

### Liver and aorta proteolysis steps

For protein precipitation, 5X volume of acetone (Fisher Scientific, catalog #A949-1) was added to the tissue homogenates^36, 37^ and then resuspended in a denaturation buffer [6.0 M urea (Sigma-Aldrich, catalog #U4884), 2.0 M thiourea (Sigma-Aldrich, catalog #T7875), 10 mM HEPES (Boston BioProducts, catalog #BBH-75-K)]. The protein amount was determined by a Pierce 660 nm Protein Assay Reagent (Thermo Fischer Scientific, catalog #22660). Proteins (10 mg per tissue preparation) were reduced in 1.0 mM dithiothreitol (DTT, Thermo Fisher Scientific, catalog #20290) and alkylated in 5.5 mM chloroacetamide (Sigma-Aldrich, catalog #C0267). Proteolysis was performed with Pierce™ Trypsin Protease (Thermo Scientific, MS Grade, catalog #90058), and the reaction was stopped with 10% Trifluoroacetic acid for HPLC (Sigma-Aldrich, catalog #302031). The peptides were desalted using Sep-Pak C18 Classic Cartridge (Waters, catalog #WAT051910) following the manufacturer’s instructions. Using a Concentrator plus complete system (Eppendorf AG, catalog #5305000304), the peptide sample was reduced to a final volume of 0.8 ml of affinity precipitation buffer (50 mM Tris-HCl pH 7.4, 10 mM MgCl_2_; Sigma-Aldrich, catalog #63069), 250 µM DTT, 50 mM NaCl]. Peptide amount was determined using a NanoDrop2000 Spectrophotometer at 280 nm (Thermo Fisher Scientific).

### Mass spectrometry for tissue proteomes

Peptides were analyzed in data-dependent acquisition (DDA) mode using the Exploris 480 fronted with an EASY-Spray Source, coupled to an Easy-nLC1200 HPLC pump (Thermo Fisher Scientific). Peptides were subjected to a dual column set-up: an Acclaim™ PepMap™ 100 C18 HPLC Column, 75 µm X 20 mm (Thermo Fisher Scientific, Cat# 164946); and an EASY-Spray™ HPLC Column, 75 µm X 250 mm (Thermo Fisher Scientific, Cat# ES902). The analytical gradient was run at 300 nl/min from 5 to 21 % Solvent B (95% acetonitrile /0.1% formic acid) for 50 minutes, followed by ten minutes of 21 to 30% Solvent B, and another 10 minutes of a sawtooth wash (alternating between 5 and 95% Solvent B) to clean the column. Solvent A was water/0.1% formic acid. The instrument was set to 120 K resolution, and the top N precursor ions in a 3-second cycle time were subjected to MS/MS (HCD collision energies (%): 24,26,28). Dynamic exclusion was enabled (60 seconds), the isolation width was 1.2 m/z, and the resolution was 120K (maximum injection time set to “auto”). Aorta and liver peptides were analyzed using a segmented survey scan, that is, three separate survey scan injections: m/z 400-1100, m/z 400-800, and m/z 700-1100.

### MS/MS spectral annotation for the tissue proteomes

The spectral processing steps were analyzed using Proteome Discoverer (version 2.5, Thermo Fisher Scientific). The spectra were queried against the Uniprot mouse (n = 63,603 entries) database (downloaded January 2022) using the SEQUEST-HT algorithm. Trypsin (full) was set as the digestion enzyme, allowing up to 2 missed cleavages and a minimum peptide length of 6 amino acids. Oxidation (+15.995 Da) of methionine and acetylation (+42.011 Da) of the N-terminus were set as variable modifications. Carbamidomethylation (+57.021 Da) of cysteine was set as a static modification. Spectral search tolerances were 10 ppm for the precursor mass and 20 mmu for HCD spectra. The peptide FDR was calculated using Percolator (target/decoy method, separate databases), and spectra were filtered based on a 1.0% FDR. Relative quantification was performed by the Feature Mapper and Precursor Ions Quantifier nodes. The maximum retention time shift for chromatographic alignments was set to 10 minutes, and the mass tolerance was set to 10 ppm. Feature linking and mapping retention time tolerance was 0, and mass tolerance was 0 ppm with a signal-to-noise threshold of 5.

### Network analysis

To generate protein-protein interaction networks that link to the Gene Ontology^38^ and Tabula Muris databases,^39^ gene identifiers were entered into the online Enrichr-KG web tool^40^ or the String Database.^41^ Each tool generates graphical representations of pathway enrichment outputs from the available Enrichr libraries.^42^

### Statistical analysis

For each tissue and timepoint proteome, we performed a two-group comparison (high-fat diet *vs*. chow diet) using the log-transformed protein group means. For each protein, we applied a Student’s t-test, and the resulting p-values were adjusted for multiple testing using the Benjamini-Hochberg procedure to control the FDR (reported as q-values).

## RESULTS

### The *Ldlr^-/-^* plasma proteome

In this study, blood was drawn from the left ventricle of *Ldlr^-/-^* mice, from which we could allocate 75 µl of plasma for nanoparticle-dependent proteomics. We analyzed plasma at 3-and 6-months of chow diet or HFD feeding, with 27 mice per diet regimen for the 3-month group and 12 mice per diet regimen for the 6-month group. We also aliquoted an additional 5 µl of plasma from 8 mice (3-month group) for a standard proteolysis workflow, referred to as the direct digestion method. The tryptic peptides from each plasma preparation were analyzed on the Orbitrap Astral using data-independent acquisition mass spectrometry (Figure 1A).

**Figure 1.**
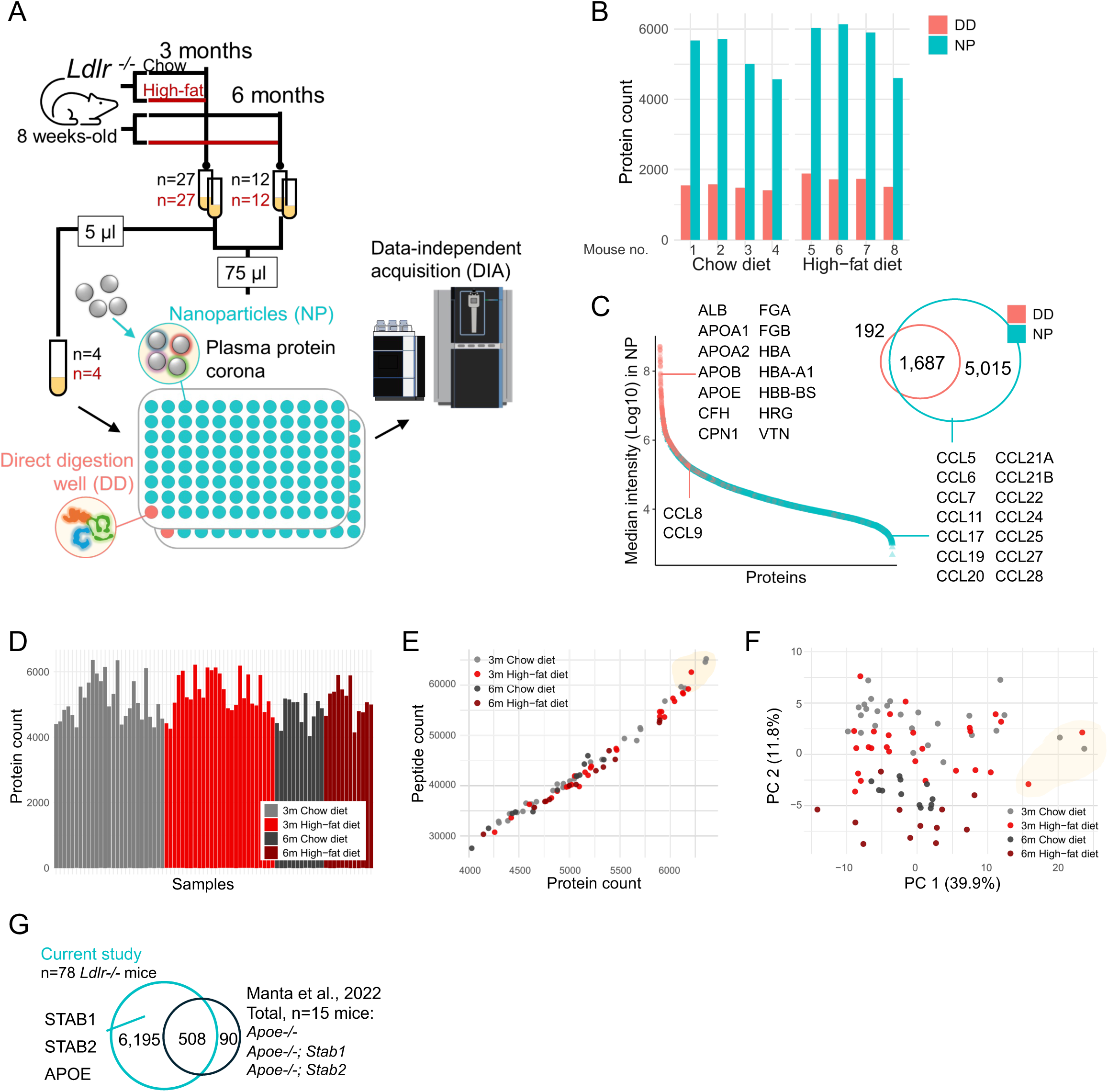
Nanoparticle-dependent deep mouse plasma proteomics (Seer’s Proteograph® platform). **(A)** Blood/plasma collection from *Ldlr^-/-^* mice fed a chow or high-fat diet for 3 (n = 27 mice/plasma samples per diet) or 6 months (n = 12 mice per diet). Eight samples were also processed using a conventional direct digestion protocol. All peptides were analyzed using the Orbitrap Astral in DIA mode. **(B, C)** A comparison between the 8 direct digest samples (8 distinct mice plasma) and their corresponding nanoparticle preparations for protein count **(B)** and median intensity **(C)** - the proteins identified in the direct digestions were mapped to their corresponding ranks in the nanoparticle data. **(D)** Protein identifications across the 78 plasma mice plasma samples. **(E)** Correlation between peptide and protein count (n = 78 samples). **(F)** Principal component analysis (n = 78) highlighting in yellow that the four distinct samples in PC-1 correspond to the samples with the most protein identifications **(E)**. **(G)** Overlap in two independent murine plasma proteome studies.

We first present the comparisons between the nanoparticle and direct digestion methods prepared from the same eight *Ldlr^-/-^* mice (Figure 1B). The direct digestion method yielded between 1,409 and 1,733 proteins (including single peptide hits with an FDR > 1%; DIA-NN), whereas the nanoparticles improved proteome depth substantially, achieving between 4,028 to 6,360 proteins across the eight mice; resulting in an increase ranging from 3- to 3.7-fold relative to the direct digestion method (Figure 1B). The proteins captured from the direct digests map to the most abundant proteins from the nanoparticle method (Figure 1C). When considering all 78 plasma samples, over 7,000 proteins were identified, and despite being noticeably lipid-laden upon collection and preparation, peptide and protein yields from the HFD plasma samples were comparable to those from chow (Figure 1D, E). A few plasma samples yielded an excess of 6,000 proteins accounting for the most variation in the data (Figure 1F, PC-1); otherwise, the 3- and 6-month exhibited visual separation in the principal component analysis (Figure 1F, PC-2). To underscore the advancement in deep mouse plasma proteome profiling, we compared our data with a recent study that employed a magnetic carboxylate-modified bead-enrichment strategy to profile the consensus plasma proteome of three genotypes, including *Apoe^-/-^*.^43^ Despite differences in mouse strains, experimental design, and workflows, 85% (508 out of 598) of the proteins identified by Manta and colleagues were also detected in our data. Unique to our dataset, however, are STAB1 and STAB2, the proteins of interest that were not captured in the previous plasma proteomes (Figure 1G).

### Diet and time-dependent changes to the *Ldlr^-/-^* plasma proteome

We next filtered the data to exclude proteins that represent a minority of mice (Figure 2A) by applying a 70% completeness threshold^44^ for either diet group per timepoint comparison; that is, including proteins identified in 19/27 for the 3-month diet comparison and 8/12 for the 6-month comparison. The filtered proteome was reduced to 5,080 proteins with 4,858 and 4,691 proteins for the 3- and 6-month groups, respectively (Figure 2B).

**Figure 2.**
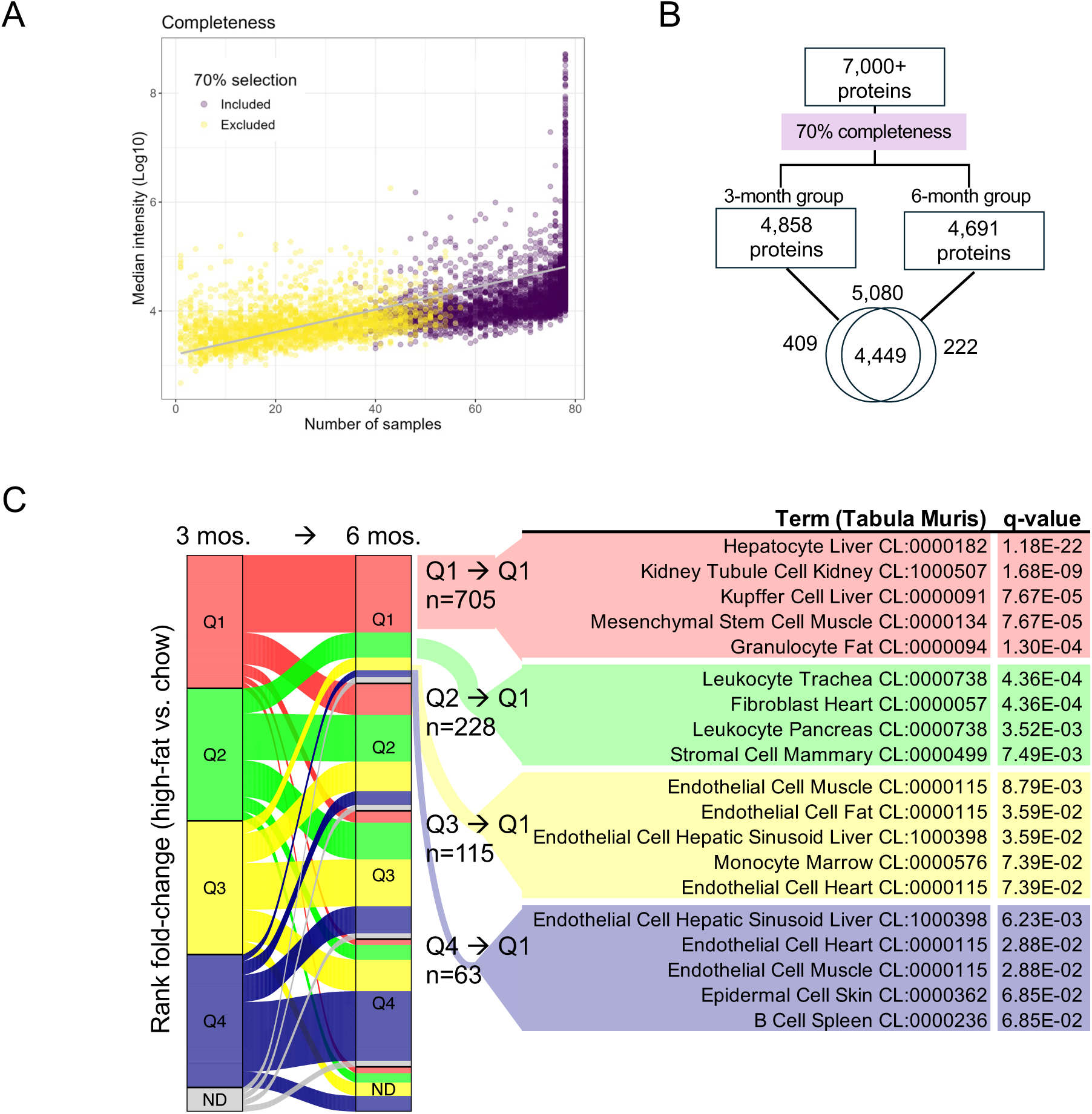
Diet and time–dependent changes to the *Ldlr*^-/-^ plasma proteome. **(A)** Protein (median intensities) plotted by the number of samples in which they were identified. Proteins included in 70% of samples (per 3-month and 6-month diet comparison) are likelier to have high-intensity signals. **(B)** Summary of the final protein sets analyzed in this study, filtered for 70% completeness in each 3-month and 6-month groups. **(C)** Alluvial plot depicting the 3-to-6-month inter-quartile relationships of proteins arranged by fold-change (high-fat vs. chow diet). The Tabula Muris single-cell database was queried using proteins from the featured alluvia, and the top 5 cell/tissues are shown.

We first overviewed sweeping HFD and timepoint-dependent changes to the *Ldlr^-/-^* plasma proteome. We computed the rank of each protein based on its fold-change in response to HFD (relative to chow diet) at 3 and 6 months. Proteins were then aggregated according to their fold-change quartile. To visualize transitions between quartiles over time, we used an alluvial plot (Figure 2C). We focused on proteins whose ranks remained within the top 25th percentile at both timepoints (Q1), as well as those that migrated from the remaining quartiles at 3-month to Q1 at 6-month (Figure 2C). We extracted the corresponding proteins and queried them into the Tabula Muris single-cell atlas database,^39^ to gain insight into potential source organs or cells. Proteins with a sustained elevation in HFD compared to chow (704 proteins from 3-month Q1 to 6-month Q1, (Figure 2C), associated with markers of liver (hepatocyte and Kupffer cells) but also kidney, mesenchymal stem cells, and fat granulocytes (Figure 2C). The prevalence of liver cells in the top-ranking quartile for both timepoints is consistent with the liver as a primary source of circulating molecules. The remaining interquartile migrations indicate other source cells and organs. The Q2 to Q1 migration (228 proteins) associates with leukocytes, fibroblasts, and stromal cells, and the Q3 to Q1 (115 proteins) and Q4 to Q1 (63) migrations associate primarily with endothelial cells and monocytes (Figure 2C). Our data indicate that with high-fat feeding, proteins associated with immunity and endothelial biology become increasingly prevalent in circulation. Proteins not detected (ND) at 3 months but present in Q1 at 6 months (57), did not yield any significant Tabula Muris hits.

### Peroxisomal and mitochondrial-associated proteins are enriched in high-fat diet *Ldlr^-/-^* plasma

We then evaluated the effects of diet by comparing the means of each 3-month (27 mice per diet) and 6-month (12 mice per diet) group (Figure 3A, B). At 3 months on a HFD, 86 proteins increased and 20 decreased; at 6 months, 146 proteins increased and 49 decreased. Proteins were considered differentially abundant based on a log_2_ fold-change greater than 1 or lower than -1, and an adjusted p-value less than 0.05. Two proteins that stand out are macrophage metalloelastase-12 (MMP-12) that increased in HFD at both timepoints, and Fetuin-B, a protease inhibitor, that decreased at both timepoints. Peroxisomal 3-ketoacyl-CoA thiolases A and B (ACAA1a/b) and procollagen galactosyltransferase 1 (COLGALT1) represent proteins unique to each 3- and 6-month filters (Figure 3A, B). Of the increased proteins, 51 (28%) are common to both timepoints, including MMP-12, peroxisomal acyl-CoA oxidase (ACOX1), mitochondrially encoded cytochrome c oxidase II (MTCO2), or the mitochondrial matrix enzyme glutamate dehydrogenase 1 (GLUD1). This overlap is encouraging, especially considering that the 3-month and 6-month plasma samples were not paired; the mice came from separate experiments, as each time point involved a terminal sacrifice.

**Figure 3.**
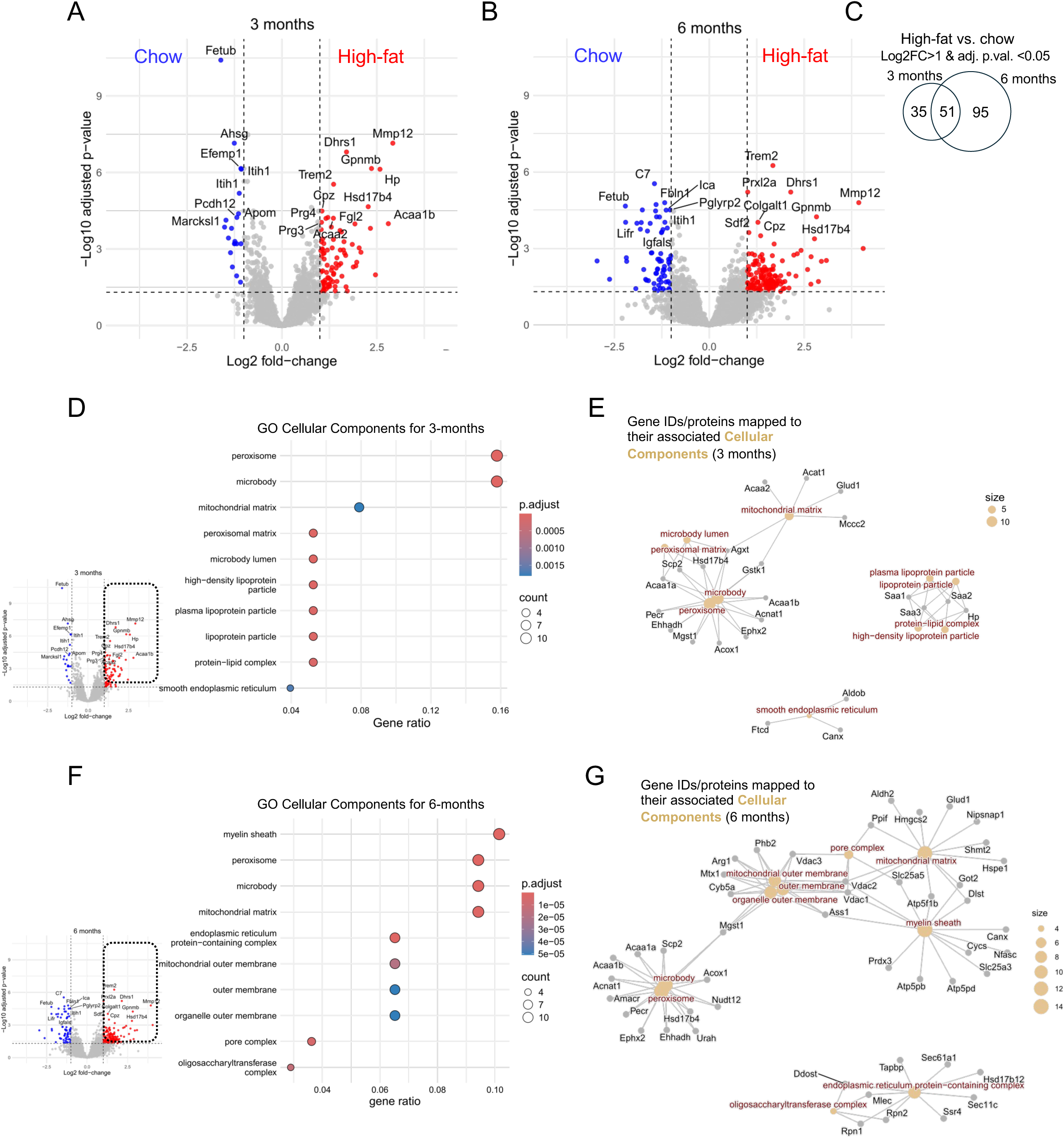
High-fat diet-induced changes to the *Ldlr*^-/-^ proteome and associated cellular components. **(A, B)** Volcano plots depicting the group mean comparisons (i.e., mean high-fat diet vs mean chow diet) for each 3-month and 6-month timepoint. Red and blue points indicate proteins with log2 fold-change > 1 (red) or < 1 (blue) with an adjusted p-value (q < 0.05). n = 27 mice per 3-month group, and n = 12 mice for each 6-month group. (C) A comparison of the uniquely and commonly identified high-fat diet-associated protein abundances from *A* and *B*. **(D, F)** Gene Ontology (GO) analyses for the high-fat diet-associated increases in the *Ldlr-/-* proteome at 3-month **(D)** and 6-month **(F)**. Miniature volcano plots are from panels **(A** and **B)**. The top-10 cellular components are listed. **(E, G)** Corresponding networks depicting the protein/gene identifiers (IDs) to cellular component associations for **(E)** and **(F)**.

We examined more closely the top-ranking HFD-associated proteins (log_2_ fold-change larger than 1 and adjusted p-value smaller than 0.05) for each timepoint using the Gene Ontology (GO) Cellular Components database (Figure 3D-G). At 3 months, the top 10 cellular components retrieved from our protein lists associated primarily with mitochondrial and peroxisomal compartments, but also lipoproteins (Figure 3D, E). The lipoproteins terms are not surprising since they comprise the most abundant particle in plasma, but it is interesting to note that it is the serum amyloid proteins (SAA1, SAA2, and SAA3) that associate with this ontology (Figure 3E). Pro-inflammatory conditions increase the prevalence of HDL-associated serum amyloids, decreasing the particle’s anti-atherogenic potential.^45^ The presence and increased abundance of mitochondrial and peroxisomal proteins are also interesting since these findings demonstrate that their proteins are secreted into circulation. At 6 months, the top-10 cellular components are also associated with peroxisomes and mitochondria, but in this case, additional proteins indicative of membrane fractions are enriched (Figure 3F, G). Taken together, the Tabula Muris (Figure 2C) and GO enrichments (Figure 3D-G), HFD feeding of *Ldlr-/-* mice may promote membranous or vesicle-mediated release of peroxisomal and mitochondrial proteins.

### The apolipoproteins exhibit distinct responses to high-fat diet feeding

Approximately 100 proteins form or bind to circulating lipoproteins in mice and humans, including the serum amyloid proteins.^46, 47^ We extracted the abundance profiles of some of the classical apolipoproteins^48^ identified in our data, and arranged them from increasing to decreasing HFD-associated changes (Figure 4). SAA1, SAA2, and SAA3 increased with HFD at 3 and 6 months of feeding, but only SAA3 surpassed the statistical cut-off (adj. p-value < 0.05) at both timepoints (Figure 4). Of the remaining apolipoproteins in our analysis, only APOC2 exhibited a similar profile to SAA3 (Figure 4). On the other hand, APOE, APOA1, APOJ, LCAT (phosphatidylcholine-sterol acyltransferase), and APOM decrease with HFD, in one or both timepoints (Figure 4). APOA2, APOA4, APOA5, APOC1, APOC4, APOD, and APOB remain constant between chow and HFD (Figure 4). APOA1 is the main structural protein for HDL and LCAT is a hallmark lipid metabolizing enzyme residing on HDL.^49^ Thus, the decrease in APOA1 and LCAT, combined with the increases in the serum amyloids together may indicate a loss of normal HDL function with high-fat feeding.

**Figure 4.**
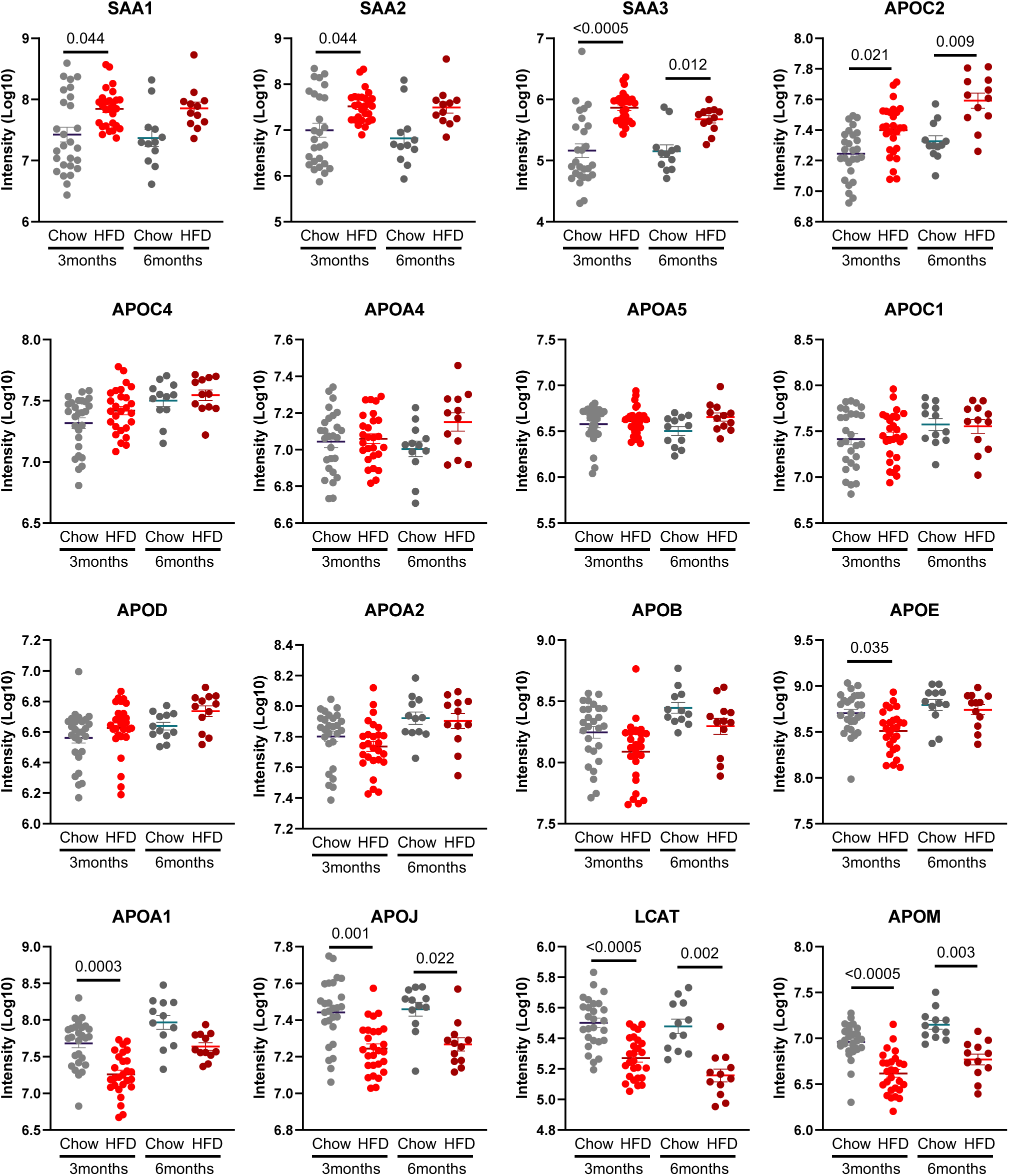
*Ldlr^-/-^* plasma *a*polipoprotein profiles. Featured apolipoproteins are ordered from increasing to decreasing abundances associated with high-fat diet feeding. T-test for each protein’s comparison and the Benjamini-Hochberg procedure to calculate the adjusted p-value. Grey, chow diet; red, high-fat diet.

### Circulating extracellular matrix components are detected in diseased aorta

Using the 3-month timepoint, we compared *Ldlr^-/-^* plasma proteome to those defining the diseased liver and aorta (i.e., fatty liver and atherosclerotic aorta). The liver and aorta samples were collected for an independent study; therefore, differences in the sample processing exist. The tissues’ peptides were analyzed using the Exploris 480 in data-dependent acquisition mode. Ten aortas were pooled for each biological replicate (n = 4 replicates) per diet. Meanwhile, liver samples were acquired from six independent mice per diet. The aorta proteome (2 or more unique peptides; chow and HFD combined) comprises 1,548 proteins, and the liver proteome, 2,142 proteins. While thirty-three percent of the plasma proteome (1,622/4,858 total plasma proteins) was shared with liver and/or aorta, plasma shared more proteins with the liver than the aorta (Figure 5A). We used GO analysis via the String Database to summarize the top 5 biological processes associated with these overlapping proteins. For proteins common to plasma and aorta, most of them are associated with extracellular matrix functions, namely the collagens and MMPs (Figures 5B, C). Proteins common to plasma and liver are associated with metabolic processes related to amino acids and other acids (Figure 5D), and proteins common to all tissues are associated with respiration pathways (Figure 5E).

**Figure 5.**
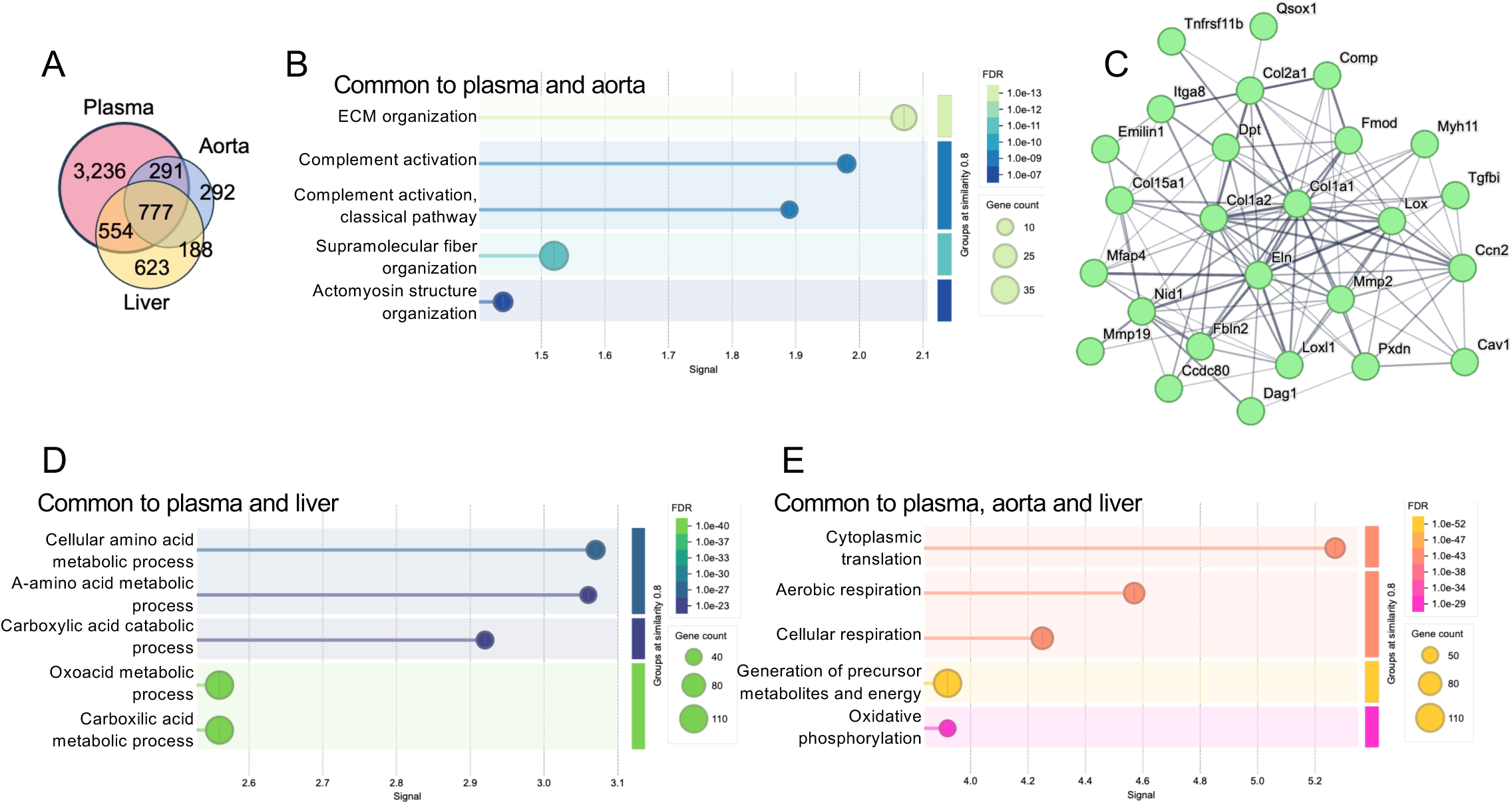
Inter-tissue comparisons using the String Database. **(A)** Venn diagram comparing *Ldlr^-/-^* plasma, liver, and aorta proteomes. Liver and aorta were acquired using DDA performed on an Orbitrap Exploris 480. **(B)** The genes comprising the top 5 Gene Ontology Biological Process common to plasma and aorta proteomes. **(C)** Protein-protein interaction network for the proteins in ECM organization Biological Process in panel *B***. (D, E)** The top 5 Biological Processes for the proteins defining plasma and liver overlap **(D)**, and plasma, aorta, and liver overlap **(E)**.

### Genes/proteins implicated in CAD are detected in *Ldlr^-/-^* plasma

We know that disease progression in *Ldlr^-/-^* mice does not fully reflect the human condition,^8^ but preclinical models are nonetheless essential for developing and testing candidate therapies. Human genome-wide association studies (GWAS) have implicated several candidate genes in CAD.^50^ We accessed a list of these CAD-associated genes curated by Aherrahrou and colleagues^50^ and confirmed that 120 of the 419 genetic products/proteins were detected in the *Ldlr^-/-^* plasma proteome (Figure 6A). Of these 120 proteins, 24 were detected in our inter-quartile range analysis (Figure 2C) and 4 exhibited marked increase in abundance with HFD, at one or two timepoints: lipoprotein lipase (LPL), very-long-chain 3-oxoacyl-CoA reductase (HSD17B12), and serine protease HTRA1 increased at 6 months whereas collagen alpha-3(VI) chain (COL6A3) at both timepoints (adj. p-value<0.05; Figure 6B, C). Whether the plasma profiles of these 4 or the remaining 116 proteins provide insight or meaning into the human condition is currently unknown. However, the ability to detect them in *Ldlr^-/-^* mouse plasma positions us to monitor them in future preclinical studies.

**Figure 6.**
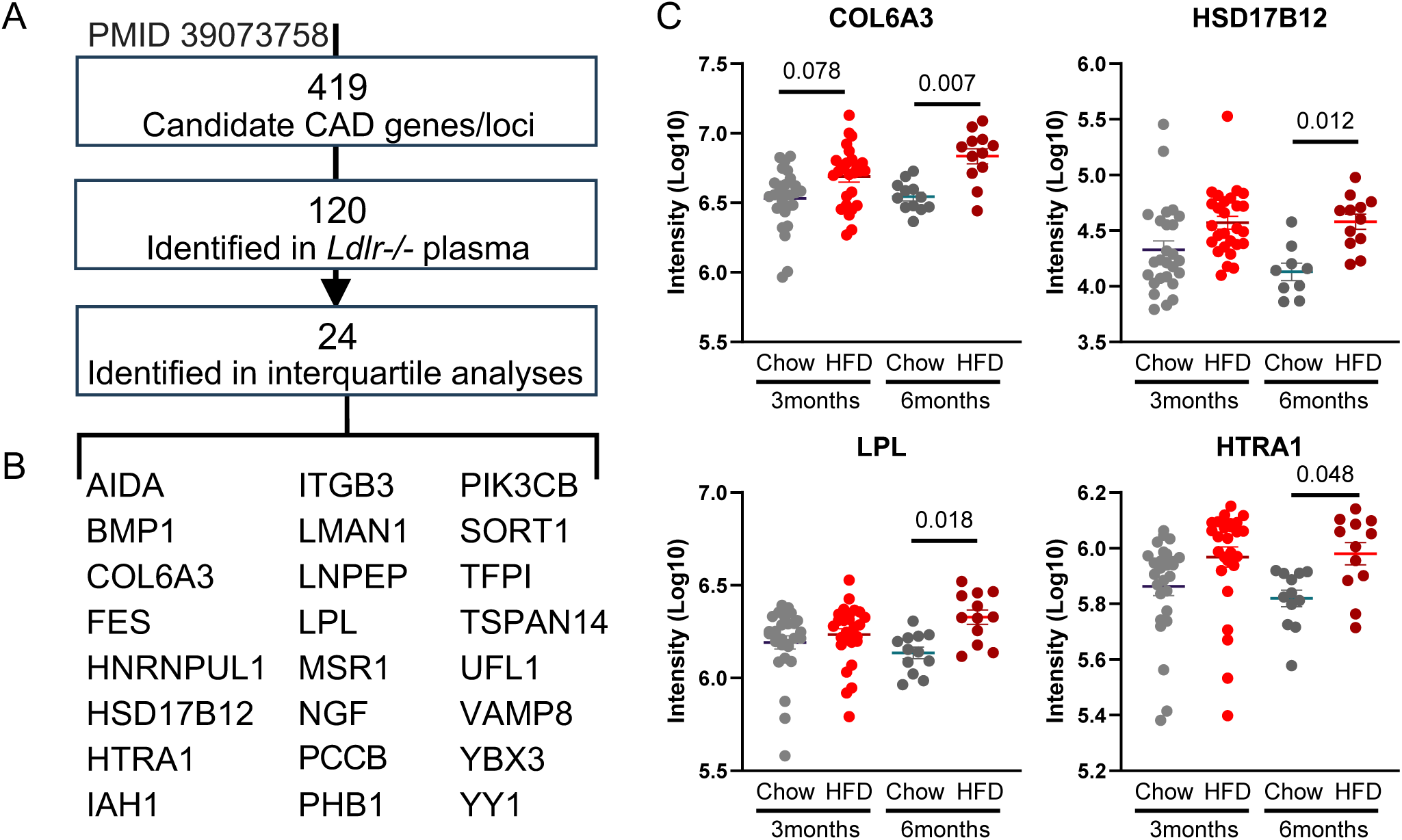
Candidate CAD-associated genes/proteins identified in *Ldlr^-^*^/-^ plasma. **(A)** List of candidate CAD-associated loci/genes curated by Aherrahrou et al. (2024), cross-referenced with the proteins identified in our study. 24 proteins were extracted from the interquartile analysis in Figure 2. **(C)** Proteins that increased in high-fat diet (HFD) over chow in one or both timepoints. T-test for each protein’s comparison and the Benjamini-Hochberg procedure to calculate the adjusted p-value.

## DISCUSSION

This study presents the most comprehensive plasma proteome of an experimental atherosclerosis model, specifically the *Ldlr*^-/-^ mouse model, a widely used preclinical species for studying the pathogenesis of atherosclerosis and metabolic disorders. High-fat feeding for three to six months in eight to ten-week-old *Ldlr*^-/-^ mice was sufficient to induce aortic lesions, as previously demonstrated.^13, 18, 29, 30^ Therefore, we examined the plasma profile for signatures that could potentially reflect disease progression. We quantified over 5,000 proteins representing 78 *Ldlr*^-/-^ mice analyzed at either 3 or 6 months of chow or HFD feeding, surpassing a previous *Apoe*^-/-^ plasma report by 10-fold.^43^

To conduct deep plasma proteomics, we relied on recently developed nanoparticle-dependent protein corona-forming technologies that have only been reported for mostly human studies.^23^ In addition, the recent advent of the Astral mass analyzer that, when implemented as a hybrid mass spectrometer, provides both fast and sensitive performance scans optimal for unprecedented comprehensive proteomics.^51, 52^ Only 75 microliters of plasma (used for two assays per mouse) were sufficient for profiling. We isolated plasma from terminal blood draws of 1,000 to 2,000 µl; however, non-terminal studies that rely on tail vein or retrobulbar venous sinus bleeding can draw up to 250 µl of blood, which is sufficient for our workflow.

The marked increase in MMP-12 at both 3 and 6 months of high-fat feeding is particularly striking. Plasma MMP-12 levels are higher in individuals with progressive atherosclerosis, aortic valve disease, and abdominal aortic aneurysm, ^53–58^ implicating MMP-12 as a potential biomarker for vessel disease progression in at least the *Ldlr*^-/-^ mice. Further mechanistic studies could elucidate whether MMP-12 directly contributes to plaque instability or merely reflects inflammatory remodeling, thus refining its utility as a therapeutic target or biomarker.

Apolipoproteins revealed distinct patterns of response to HFD feeding in *Ldlr*^-/-^ mice, reflecting the complex alterations in lipoprotein metabolism during atherogenesis. The observed decrease in APOA1 and LCAT is notable given their pivotal roles in HDL particle formation and cholesterol efflux. APOA1 is the major structural protein of HDL, and LCAT is critical for HDL maturation and reverse cholesterol transport; reductions in these proteins likely impair HDL’s atheroprotective functions, consistent with prior reports that HDL dysfunction accompanies CAD.^59–61^ Conversely, the increase in plasma serum amyloid proteins SAA1-3 further suggests a shift towards a pro-inflammatory HDL phenotype, known to reduce HDL’s anti-inflammatory and antioxidant properties.^62^ Of note, APOC2, a cofactor for lipoprotein lipase LPL, also increased with HFD, potentially indicating an adaptive response to elevated triglyceride-rich lipoproteins. These dynamic changes underscore the importance of apolipoprotein remodeling in the pathogenesis of diet-induced atherosclerosis.

The hepatokine fetuin-B, a member of the superfamily of cysteine protease inhibitors, decreased in plasma with HFD. Fetuin-B is involved in metabolic regulation and inhibition of ectopic calcification, but its role in the cardiovascular context remains less clear that that of fetuin-A.^63^ Decreased circulating fetuin-B levels could reflect hepatic dysfunction or altered systemic inflammatory states associated with diet-induced atherosclerosis. Recent studies suggest fetuin-B may influence insulin resistance and lipid metabolism,^64^ linking it to metabolic pathways that intersect with atherogenesis.^65^ Our data position fetuin-B as a novel candidate for further exploration and a possible modulator of metabolic and inflammatory aspects of cardiovascular disease.

The comparative analysis of the plasma, aorta, and liver proteomes revealed distinct yet interrelated molecular signatures that reflect the systemic and tissue-specific impact of diet-induced atherosclerosis. The plasma proteome captured not only circulating inflammatory and metabolic proteins but also key signals indicative of underlying tissue pathology. Notably, proteins involved in fatty acid beta-oxidation and lipid metabolism were enriched in both liver and plasma, highlighting the liver’s central role in metabolic dysregulation during HFD feeding and its systemic repercussions. Conversely, the aortic proteome was characterized by elevated extracellular matrix remodeling proteins and inflammatory mediators, consistent with vascular lesion formation and progression. The overlap of certain plasma proteins with those in the aorta suggests that circulating proteins may serve as accessible biomarkers reflecting vascular remodeling processes. This inter-tissue proteomic concordance supports the concept that plasma can function as a window into pathophysiological changes occurring within specific organs. These findings advocate for the utility of combining plasma proteomics with tissue profiling to improve biomarker discovery and to elucidate the crosstalk between metabolic and vascular compartments in atherosclerosis.

However, despite these advances, limitations remain. The *Ldlr*^-/-^ mouse, while highly informative, may not fully recapitulate human atherosclerosis complexity, and species-specific differences in proteomic profiles warrant careful interpretation, as do the potential differences owing to the animal model (e.g., *Apoe^-/-^* vs. *Ldlr^-/-^).* We also anticipate that deep plasma proteomic efforts will lead to proteoform discovery,^27, 66^ that is, the types of posttranslational modified proteins that circulate in these mice. Plasma proteoform profiling will incite new technologies for enrichment and new avenues for biomarker discovery. Future studies incorporating complementary omics approaches, such as lipidomics and transcriptomics, may enhance biomarker discovery and mechanistic insight.

In conclusion, we demonstrate that nanoparticle-dependent enrichment of low-abundant plasma proteins coupled to mass spectrometry for proteome sequencing offers a viable and powerful strategy to monitor disease-relevant molecular signals in a mouse model of cardiometabolic disorders (e.g., dyslipidemia, atherosclerosis and fatty liver). This study lays the groundwork for future efforts to assess the predictive value of circulating proteins in atherosclerosis and to test whether they can serve as readouts for therapeutic efficacy in preclinical intervention studies. Ultimately, such proteomic strategies could accelerate the translation of preclinical findings to clinical diagnostics and for cardiovascular disease.

## ABBREVIATIONS

*Apoe^-/-^*: Apolipoprotein E-deficient mice
CAD: Coronary artery disease
CHD: Coronary heart disease
DDA: Data-dependent acquisition
DIA: Data-independent acquisition
FDR: False-discovery rate
GWAS: Genome-wide association study
HDL: High-density lipoproteins
HFD: High-fat diet
IDL: Intermediate-density lipoprotein
LDL: Low-density lipoprotein
LDL-C: Low-density lipoprotein cholesterol
LDLR: Low-density lipoprotein receptor
*Ldlr^-/-^*: Low-density lipoprotein receptor-deficient mice
VLDL: Very low-density lipoprotein

## Reference

1. Martin SS, Aday AW, Almarzooq ZI, et al. 2024 Heart Disease and Stroke Statistics: A Report of US and Global Data From the American Heart Association. Circulation. 2024;149:e347–e913.

2. Watkin DM, Lawry EY, Mann GV and Halperin M. A study of serum beta lipoprotein and total cholesterol variability and its relation to age and serum level in adult human subjects. J Clin Invest. 1954;33:874–83.

3. Barr DP, Russ EM and Eder HA. Protein-lipid relationships in human plasma. II. In atherosclerosis and related conditions. Am J Med. 1951;11:480–93.

4. Castelli WP, Cooper GR, Doyle JT, et al. Distribution of triglyceride and total, LDL and HDL cholesterol in several populations: a cooperative lipoprotein phenotyping study. J Chronic Dis. 1977;30:147–69.

5. Castelli WP, Doyle JT, Gordon T, Hames CG, Hjortland MC, Hulley SB, Kagan A and Zukel WJ. HDL cholesterol and other lipids in coronary heart disease. The cooperative lipoprotein phenotyping study. Circulation. 1977;55:767–72.

6. Gofman JW, Young W and Tandy R. Ischemic heart disease, atherosclerosis, and longevity. Circulation. 1966;34:679–97.

7. Getz GS and Reardon CA. Animal models of atherosclerosis. Arterioscler Thromb Vasc Biol. 2012;32:1104–15.

8. Getz GS and Reardon CA. Do the Apoe-/- and Ldlr-/- Mice Yield the Same Insight on Atherogenesis? Arterioscler Thromb Vasc Biol. 2016;36:1734–41.

9. Plump AS, Smith JD, Hayek T, Aalto-Setala K, Walsh A, Verstuyft JG, Rubin EM and Breslow JL. Severe hypercholesterolemia and atherosclerosis in apolipoprotein E-deficient mice created by homologous recombination in ES cells. Cell. 1992;71:343–53.

10. Ishibashi S, Brown MS, Goldstein JL, Gerard RD, Hammer RE and Herz J. Hypercholesterolemia in low density lipoprotein receptor knockout mice and its reversal by adenovirus-mediated gene delivery. J Clin Invest. 1993;92:883–93.

11. Zhang SH, Reddick RL, Piedrahita JA and Maeda N. Spontaneous hypercholesterolemia and arterial lesions in mice lacking apolipoprotein E. Science. 1992;258:468–71.

12. Hartvigsen K, Binder CJ, Hansen LF, et al. A diet-induced hypercholesterolemic murine model to study atherogenesis without obesity and metabolic syndrome. Arterioscler Thromb Vasc Biol. 2007;27:878–85.

13. Getz GS and Reardon CA. Diet and murine atherosclerosis. Arterioscler Thromb Vasc Biol. 2006;26:242–9.

14. Tabas I, Williams KJ and Boren J. Subendothelial lipoprotein retention as the initiating process in atherosclerosis: update and therapeutic implications. Circulation. 2007;116:1832–44.

15. Bouchareb R, Boulanger MC, Tastet L, et al. Activated platelets promote an osteogenic programme and the progression of calcific aortic valve stenosis. Eur Heart J. 2019;40:1362–1373.

16. Chaix A, Lin T, Ramms B, et al. Time-Restricted Feeding Reduces Atherosclerosis in LDLR KO Mice but Not in ApoE Knockout Mice. Arterioscler Thromb Vasc Biol. 2024;44:2069–2087.

17. Morgan S, Lee LH, Halu A, et al. Identifying novel mechanisms of abdominal aortic aneurysm via unbiased proteomics and systems biology. Front Cardiovasc Med. 2022;9:889994.

18. Yang ZH, Gordon SM, Sviridov D, et al. Dietary supplementation with long-chain monounsaturated fatty acid isomers decreases atherosclerosis and alters lipoprotein proteomes in LDLr(-/-) mice. Atherosclerosis. 2017;262:31–38.

19. Amor M, Bianco V, Buerger M, et al. Genetic deletion of MMP12 ameliorates cardiometabolic disease by improving insulin sensitivity, systemic inflammation, and atherosclerotic features in mice. Cardiovasc Diabetol. 2023;22:327.

20. Geyer PE, Kulak NA, Pichler G, Holdt LM, Teupser D and Mann M. Plasma Proteome Profiling to Assess Human Health and Disease. Cell Syst. 2016;2:185–95.

21. Geyer PE, Holdt LM, Teupser D and Mann M. Revisiting biomarker discovery by plasma proteomics. Mol Syst Biol. 2017;13:942.

22. Singh B and Mayr M. Enhancing cardiovascular risk prediction through proteomics? Cardiovasc Res. 2024;120:e2–e4.

23. Ferdosi S, Tangeysh B, Brown TR, et al. Engineered nanoparticles enable deep proteomics studies at scale by leveraging tunable nano-bio interactions. Proc Natl Acad Sci U S A. 2022;119:e2106053119.

24. Vroman L, Adams AL, Fischer GC and Munoz PC. Interaction of high molecular weight kininogen, factor XII, and fibrinogen in plasma at interfaces. Blood. 1980;55:156–9.

25. Ferdosi S, Stukalov A, Hasan M, et al. Enhanced Competition at the Nano-Bio Interface Enables Comprehensive Characterization of Protein Corona Dynamics and Deep Coverage of Proteomes. Adv Mater. 2022;34:e2206008.

26. Kverneland AH, Harking F, Vej-Nielsen JM, Huusfeldt M, Bekker-Jensen DB, Svane IM, Bache N and Olsen JV. Fully Automated Workflow for Integrated Sample Digestion and Evotip Loading Enabling High-Throughput Clinical Proteomics. Mol Cell Proteomics. 2024;23:100790.

27. Huang CF, Hollas MA, Sanchez A, et al. Deep Profiling of Plasma Proteoforms with Engineered Nanoparticles for Top-Down Proteomics. J Proteome Res. 2024;23:4694–4703.

28. Suhre K, Venkataraman GR, Guturu H, et al. Nanoparticle enrichment mass-spectrometry proteomics identifies protein-altering variants for precise pQTL mapping. Nat Commun. 2024;15:989.

29. Olkowicz M, Tomczyk M, Debski J, et al. Enhanced cardiac hypoxic injury in atherogenic dyslipidaemia results from alterations in the energy metabolism pattern. Metabolism. 2021;114:154400.

30. Ma Y, Wang W, Zhang J, Lu Y, Wu W, Yan H and Wang Y. Hyperlipidemia and atherosclerotic lesion development in Ldlr-deficient mice on a long-term high-fat diet. PLoS One. 2012;7:e35835.

31. Parasuraman S, Raveendran R and Kesavan R. Blood sample collection in small laboratory animals. J Pharmacol Pharmacother. 2010;1:87–93.

32. Blume JE, Manning WC, Troiano G, et al. Rapid, deep and precise profiling of the plasma proteome with multi-nanoparticle protein corona. Nat Commun. 2020;11:3662.

33. Demichev V, Messner CB, Vernardis SI, Lilley KS and Ralser M. DIA-NN: neural networks and interference correction enable deep proteome coverage in high throughput. Nat Methods. 2020;17:41–44.

34. Olinger B, Banarjee R, Dey A, et al. The secretome of senescent monocytes predicts age-related clinical outcomes in humans. Nat Aging. 2025.

35. Kuraoka S, Higashi H, Yanagihara Y, et al. A Novel Spectral Annotation Strategy Streamlines Reporting of Mono-ADP-ribosylated Peptides Derived from Mouse Liver and Spleen in Response to IFN-gamma. Mol Cell Proteomics. 2022;21:100153.

36. Larsen SC, Leutert M, Bilan V, Martello R, Jungmichel S, Young C, Hottiger MO and Nielsen ML. Proteome-Wide Identification of In Vivo ADP-Ribose Acceptor Sites by Liquid Chromatography-Tandem Mass Spectrometry. Methods Mol Biol. 2017;1608:149–162.

37. Higashi H, Maejima T, Lee LH, Yamazaki Y, Hottiger MO, Singh SA and Aikawa M. A Study into the ADP-Ribosylome of IFN-gamma-Stimulated THP-1 Human Macrophage-like Cells Identifies ARTD8/PARP14 and ARTD9/PARP9 ADP-Ribosylation. J Proteome Res. 2019;18:1607–1622.

38. Gene Ontology C. Gene Ontology Consortium: going forward. Nucleic Acids Res. 2015;43:D1049–56.

39. Tabula Muris C, Overall c, Logistical c, et al. Single-cell transcriptomics of 20 mouse organs creates a Tabula Muris. Nature. 2018;562:367–372.

40. Evangelista JE, Xie Z, Marino GB, Nguyen N, Clarke DJB and Ma’ayan A. Enrichr-KG: bridging enrichment analysis across multiple libraries. Nucleic Acids Res. 2023;51:W168–W179.

41. Konjevoda M. [Successful pregnancies after endometrial biopsies during the conception cycle]. Jugosl Ginekol Opstet. 1982;22:146–8.

42. Chen EY, Tan CM, Kou Y, Duan Q, Wang Z, Meirelles GV, Clark NR and Ma’ayan A. Enrichr: interactive and collaborative HTML5 gene list enrichment analysis tool. BMC Bioinformatics. 2013;14:128.

43. Manta CP, Leibing T, Friedrich M, et al. Targeting of Scavenger Receptors Stabilin-1 and Stabilin-2 Ameliorates Atherosclerosis by a Plasma Proteome Switch Mediating Monocyte/Macrophage Suppression. Circulation. 2022;146:1783–1799.

44. Tyanova S, Temu T, Sinitcyn P, Carlson A, Hein MY, Geiger T, Mann M and Cox J. The Perseus computational platform for comprehensive analysis of (prote)omics data. Nat Methods. 2016;13:731–40.

45. Van Lenten BJ, Hama SY, de Beer FC, Stafforini DM, McIntyre TM, Prescott SM, La Du BN, Fogelman AM and Navab M. Anti-inflammatory HDL becomes pro-inflammatory during the acute phase response. Loss of protective effect of HDL against LDL oxidation in aortic wall cell cocultures. J Clin Invest. 1995;96:2758–67.

46. Pamir N, Hutchins P, Ronsein G, Vaisar T, Reardon CA, Getz GS, Lusis AJ and Heinecke JW. Proteomic analysis of HDL from inbred mouse strains implicates APOE associated with HDL in reduced cholesterol efflux capacity via the ABCA1 pathway. J Lipid Res. 2016;57:246–57.

47. Singh SA, Andraski AB, Pieper B, Goh W, Mendivil CO, Sacks FM and Aikawa M. Multiple apolipoprotein kinetics measured in human HDL by high-resolution/accurate mass parallel reaction monitoring. J Lipid Res. 2016;57:714–28.

48. Alaupovic P. Apoliproproteins and lipoproteins. Atherosclerosis. 1971;13:141–6.

49. Glomset JA. The mechanism of the plasma cholesterol esterification reaction: plasma fatty acid transferase. Biochim Biophys Acta. 1962;65:128–35.

50. Aherrahrou R, Reinberger T, Hashmi S and Erdmann J. GWAS breakthroughs: mapping the journey from one locus to 393 significant coronary artery disease associations. Cardiovasc Res. 2024;120:1508–1530.

51. Bubis JA, Arrey TN, Damoc E, et al. Challenging the Astral mass analyzer to quantify up to 5,300 proteins per single cell at unseen accuracy to uncover cellular heterogeneity. Nat Methods. 2025;22:510–519.

52. Stewart HI, Grinfeld D, Giannakopulos A, et al. Parallelized Acquisition of Orbitrap and Astral Analyzers Enables High-Throughput Quantitative Analysis. Anal Chem. 2023;95:15656–15664.

53. Nakano T, Katsuki S, Chen M, et al. Uremic Toxin Indoxyl Sulfate Promotes Proinflammatory Macrophage Activation Via the Interplay of OATP2B1 and Dll4-Notch Signaling. Circulation. 2019;139:78–96.

54. Lim CS, Shalhoub J, Gohel MS, Shepherd AC and Davies AH. Matrix metalloproteinases in vascular disease--a potential therapeutic target? Curr Vasc Pharmacol. 2010;8:75–85.

55. Mahdessian H, Perisic Matic L, Lengquist M, et al. Integrative studies implicate matrix metalloproteinase-12 as a culprit gene for large-artery atherosclerotic stroke. J Intern Med. 2017;282:429–444.

56. Goncalves I, Bengtsson E, Colhoun HM, et al. Elevated Plasma Levels of MMP-12 Are Associated With Atherosclerotic Burden and Symptomatic Cardiovascular Disease in Subjects With Type 2 Diabetes. Arterioscler Thromb Vasc Biol. 2015;35:1723–31.

57. Curci JA, Liao S, Huffman MD, Shapiro SD and Thompson RW. Expression and localization of macrophage elastase (matrix metalloproteinase-12) in abdominal aortic aneurysms. J Clin Invest. 1998;102:1900–10.

58. Hellenthal FA, Buurman WA, Wodzig WK and Schurink GW. Biomarkers of AAA progression. Part 1: extracellular matrix degeneration. Nat Rev Cardiol. 2009;6:464–74.

59. Rosenson RS, Brewer HB, Jr., Ansell BJ, Barter P, Chapman MJ, Heinecke JW, Kontush A, Tall AR and Webb NR. Dysfunctional HDL and atherosclerotic cardiovascular disease. Nat Rev Cardiol. 2016;13:48–60.

60. Khera AV, Cuchel M, de la Llera-Moya M, et al. Cholesterol efflux capacity, high-density lipoprotein function, and atherosclerosis. N Engl J Med. 2011;364:127–35.

61. Saleheen D, Scott R, Javad S, et al. Association of HDL cholesterol efflux capacity with incident coronary heart disease events: a prospective case-control study. Lancet Diabetes Endocrinol. 2015;3:507–13.

62. Webb NR. High-Density Lipoproteins and Serum Amyloid A (SAA). Curr Atheroscler Rep. 2021;23:7.

63. Almarashda O, Abdi S, Yakout S, Khattak MNK and Al-Daghri NM. Hepatokines Fetuin-A and Fetuin-B status in obese Saudi patient with diabetes mellitus type 2. Am J Transl Res. 2022;14:3292–3302.

64. Li Z, He C, Liu Y, et al. Association of Fetuin-B with Subclinical Atherosclerosis in Obese Chinese Adults. J Atheroscler Thromb. 2020;27:418–428.

65. Wang D, Wu M, Zhang X, Li L, Lin M, Shi X, Zhao Y, Huang C and Li X. Hepatokine Fetuin B expression is regulated by leptin-STAT3 signalling and associated with leptin in obesity. Sci Rep. 2022;12:12869.

66. Donovan MKR, Huang Y, Blume JE, et al. Functionally distinct BMP1 isoforms show an opposite pattern of abundance in plasma from non-small cell lung cancer subjects and controls. PLoS One. 2023;18:e0282821.

